# Exploration-related strategy mediates negative coupling between decision-making performance and psychiatric symptoms

**DOI:** 10.1101/730614

**Authors:** Shinsuke Suzuki, Yuichi Yamashita, Kentaro Katahira

## Abstract

Various psychiatric symptoms are often accompanied by impairments in decision-making. Given the high comorbidity of psychiatric disorders, symptoms that specifically couple with the impairment remain unidentified. The computations involved in decision-making that mediate the coupling are also elusive. Here, we conducted an online experiment with healthy individuals (*n*=939), participating in a decision-making task and completing questionnaires about psychiatric symptoms. The questionnaire data identified two dimensions underlying various symptoms: the first is mainly associated with obsessive-compulsive traits whereas the second is associated with depression and anxiety. Furthermore, by model-based analysis, we found only the first dimension was negatively correlated with the overall performance of the decision-making task, and the negative correlation was mediated by choice stochasticity (random exploration) and propensity to choose options previously unchosen. Given that the preference for previously unchosen options can reflect uncertainty-driven exploration, our findings highlight a key role of exploration-related strategies in psychiatric deficits of decision-making.

---

Decision-making to attain reward is crucial for our survival, such as where to hunt (e.g., forest or lake) and where to invest (e.g., risky equities or safe bonds) ^1^. Recent studies in psychology, neuroscience, and psychiatry have suggested that impairments in decision-making are often accompanied by mental disorders such as obsessive-compulsive disorder^2,3^, depression^4^, anxiety^5^, and schizophrenia^6^. However, little is known about the mechanism that attenuates decision performance. Here in this study, we provide a mechanistic account of how certain psychiatric symptoms are associated with deficits in decision-making.

An increasing number of studies has well-described the decision-making process using a machine learning algorithm known as reinforcement learning^7^. The reinforcement learning framework posits that appropriate decisions require individuals to learn the expected value of each option from the reward experience. Furthermore, in order to maximise long-term reward, they need to choose a less familiar (more uncertain) option on some occasions in order to collect more information about its value (i.e., exploration), while choosing the sure rewarding option on most of the occasions (i.e., exploitation)^8–10^. A simple strategy of exploration is to increase choice stochasticity and this is known as random exploration^11^. A more sophisticated strategy would be driven by uncertainty about the value of each option^12^, that is, uncertain options are more likely to be chosen than promising options. The exploit-explore balance is of particular importance in changing environments. The reinforcement learning account of the decision-making process has been of considerable concern in many research fields such as psychology, economics, neuroscience, and psychiatry, as it not only captures the organisms’ behaviours but also their neural activities^13–19^.

In the last decade, by using reinforcement learning and other computational frameworks, an emerging research field, known as computational psychiatry, has attempted to uncover the types of computation underlying decision-making that is associated with psychiatric symptoms^20–23^. For example, studies suggest that imbalance between the inflexible and the flexible decision-making strategies is linked to obsessive-compulsive disorder, which is characterised by repetitive adverse behaviours (e.g., excessive hand-washing potentially leading to skin troubles)^24,25^. More specifically, individuals with high obsessive-compulsive traits tend to rely less on flexible model-based reinforcement learning^25,26^. Furthermore, depressive traits are associated with diminished sensitivity to reward feedback^27^ and the equivalent component, choice stochasticity^28^. This is consistent with the fact that anhedonia or loss of pleasure is a core symptom of major depressive disorder. Moreover, negative symptoms in schizophrenia are found to be associated with deficits in uncertainty-driven exploration in reinforcement learning^29^.

Despite the advancement in computational psychiatry, we have yet to better understand how psychiatric symptoms are coupled with deficits in decision-making. Various psychiatric symptoms are known to be accompanied by impairments in the overall performance of decision-making, and recent studies have examined associations between psychiatric symptoms and particular computational processes of decision-making^20–23^. To our knowledge, however, very few studies have provided a comprehensive view of the tripartite relationship between the particular psychiatric symptoms, overall performance of decision-making, and specific computational processes^30^. In other words, it is still unknown which computational processes mediate negative coupling between psychiatric symptoms and the performance of decision-making. Of note, the difficulty is at least in part due to a high comorbidity rate of mental disorders^26,31^. Half of the individuals with a confirmed diagnosis of one mental disorder may also have additional disorders at the same time^32^. Furthermore, correspondence between the categorical diagnosis of mental-disorder and the computational process seems to be not one-to-one. One behavioural symptom can result from several causes, and one cause can lead to various behavioural symptoms, i.e., *equifinality* and *multifinality* respectively^33,34^. The high comorbidity, multifinality, and equifinality, therefore, make it difficult to examine the relationship between particular psychiatric symptoms and decision-making.

To elucidate the tripartite relationship, we conducted a large-scale online experiment with healthy participants, an emerging tool in psychiatry useful for dealing with high comorbidity, multifinality and equifinality of mental disorders^35^. In our experiment, participants (*n* = 939) performed a reward-seeking decision-making task and then completed questionnaires about psychiatric symptoms. By employing model-based analyses together with factor and mediation analyses, we found that there are at least two dimensions underlying psychiatric symptoms; the first is mainly associated with obsessive-compulsive traits, and the second is associated with depressive and anxious traits. We then found that only the first dimension is negatively correlated with the overall performance of the decision-making task. Finally, we show that the negative correlation between the psychiatric dimension and the task performance is mediated by individual differences in choice stochasticity (i.e., random exploration) and propensity to choose options previously unchosen, which is thought to reflect uncertainty-driven exploration.

## Results

### Experimental task

In our experiment, healthy participants (*n* = 939; see Fig. S1a for demographic information) performed a reward-seeking decision-making task (i.e., conventional three-armed bandit task) and then completed questionnaires about psychiatric symptoms (schizotypal, obsessive-compulsive, depressive, anxious and impulsive traits) as well as socioeconomic status. In the decision-making task, participants repeatedly chose among three options (i.e., three fractal stimuli; see Fig. 1a). In each trial, they received a reward depending on the probability assigned to the chosen option. As the reward probability for each option was unknown to the participants and changed dynamically (Fig. 1b, *left*), participants were required to continuously learn the probabilities over the course of the trials in order to maximise reward earnings. After a careful assessment of the participants’ engagement in the experiment (see Methods), we confirmed that participants’ choice probability of each option co-varied with the change of the reward probability (Fig. 1b, *left*). Furthermore, they were more likely to choose the best option that has the highest reward probability in a given trial (*P* < 0.01; Fig. 1c), compared to the other options. These data suggest that the participants succeeded in keeping track of the changing reward probabilities.

**Fig. 1:**
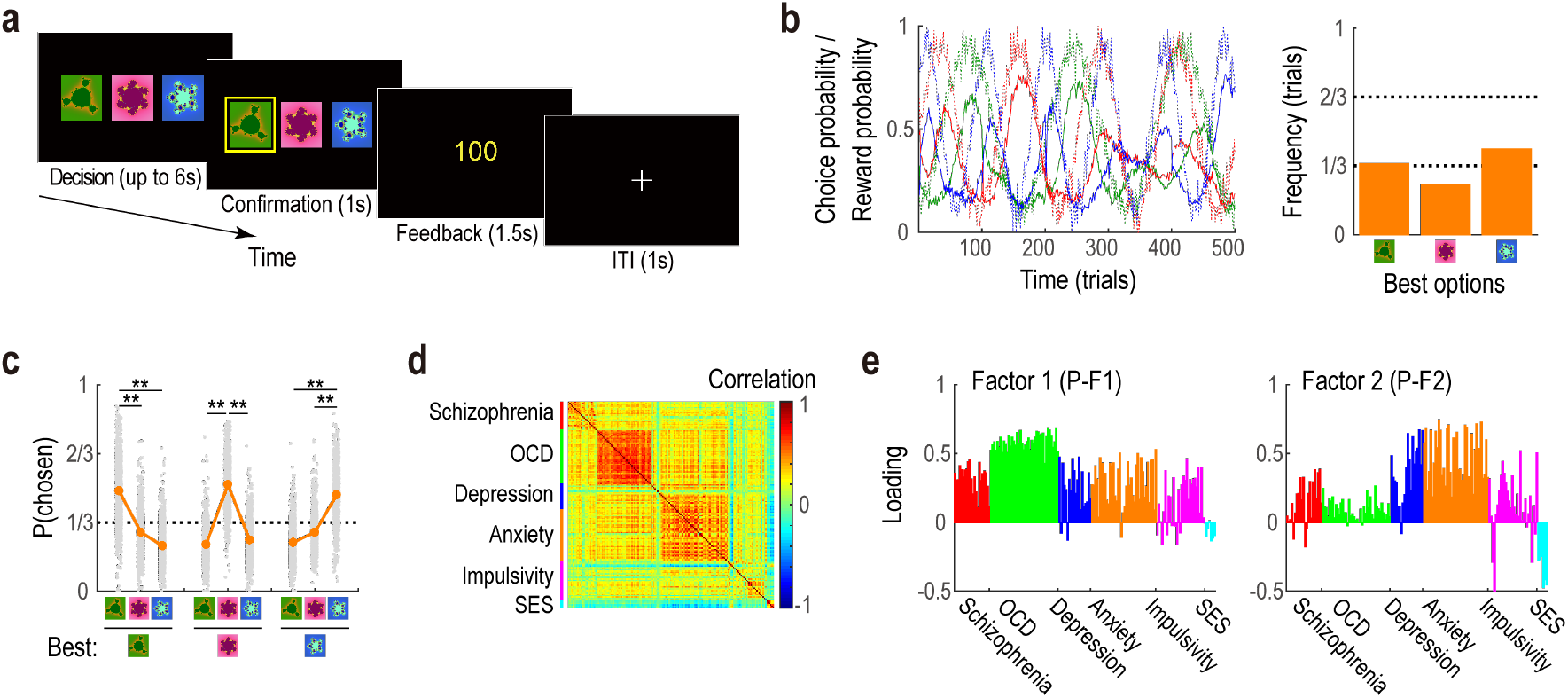
Experimental task and the questionnaire data. (a) Illustration of the reward-seeking decision-making task (i.e., three-armed bandit task). On each trial, participants chose among the three options and received a reward depending on the probability assigned to the chosen option. (b) Reward and choice probabilities for each option. Dashed and solid lines respectively depict reward probability assigned to each option and the participants’ actual choice probability of each option (left). Frequency of trials in which each of the three options had the highest reward probability (i.e., the best option) (right). (c) Proportion of each option chosen. Orange points and the error-bars denote the mean and SEM across participants (*n* = 939). Note that the error-bars are overlapped with the orange points. Each grey point indicates one participant’s data. Left panel denotes the proportions when the green option had the highest reward probability (i.e., the best option). Middle denotes the case in which the red option was the best, and right denotes the case in which the blue option was the best. \*\**P* < 0.01 (Bonferroni corrected: *α*’ = 0.01/6) in two-tailed *t*-tests (*G* vs. *R* when *G* was best: *diff* = 0.20, *t*_938_ = 25.89, *P* < 0.001; *G* vs. *B* when *G* was best: *diff* = 0.27, *t*_938_ = 37.03, *P* < 0.001; *G* vs. *R* when *R* was best: *diff* = −0.29, *t*_938_ = −43.33, *P* < 0.001; *R* vs. *B* when *R* was best: *diff* = 0.27, *t*_938_ = 39.66, *P* < 0.001; *G* vs. *B* when *B* was best: *diff* = −0.23, *t*_938_ = −35.41, *P* < 0.001; *R* vs. *B* when *B* was best: *diff* = −0.18, *t*_938_ = −27.02, *P* < 0.001). *G*, the green option; *R*, the red option; *B*, the blue option; and *diff*, a difference in the mean proportion. (d) Cross-correlation of the responses to the questionnaire items. *Red*, questionnaire items related to schizophrenia; *green*, items related to obsessive-compulsive disorder (OCD); *blue*, items related to depression; *orange*, items related to anxiety; *magenta*, items related to impulsivity; *cyan*, items related to socio-economic status (SES). (e) Loadings of the questionnaire items in the two factors underlying psychiatric symptoms. *Left*, loadings in the first factor (P-F1); and *right*, those in the second factor (P-F2). Loading values of each item are derived by the factor analysis.

### Two factors underlying psychiatric symptoms

Based on the questionnaire data, we first sought to identify the hidden factors underlying multiple psychiatric symptoms. As expected, the psychiatric symptoms measured by the questionnaires were highly associated with each other (Fig. 1d). A visual inspection of the cross-correlation matrix found that the symptoms related to obsessive-compulsive disorder, depression, and anxiety were inter-correlated (Fig. 1c). Consistent with this visual inspection, a formal factor analysis revealed at least two hidden factors underlying the psychiatric symptoms. The first factor (P-F1) has a higher loading from symptoms mainly associated with obsessive-compulsive traits (Fig. 1e, *left*) whereas the second factor (P-F2) has a higher loading from symptoms mainly associated with depression and anxiety (Fig. 1e, *right*). While we determined the number of factors based on a previous large-scale longitudinal study on patients^31^ and slopes of the scree-plot (Fig. S1b) (see Methods for details), this composition did not change even when we assumed three factors, instead of two, underlying the symptoms (Fig. S1c). Furthermore, we replicated this finding in an independent sample (*n* = 961; Fig. S1d-f). These results together support the existence of at least two dimensions underlying the mutually related psychiatric symptoms. P-F1 is mainly associated with obsessive-compulsive traits while P-F2 is associated with depression and anxiety.

### Psychiatric symptoms and overall task performance

Having established the two factors underlying the psychiatric symptoms, we next examined how these factors relate to the overall performance of the decision-making task. Linear regression analyses found that the first psychiatric factor (P-F1) had a negative impact on task performance (Fig. 2ab). The proportion of the optimal choices (i.e., choices of the best option) was negatively correlated with P-F1 across the participants (*P* < 0.01; Fig. 2a). Furthermore, the amount of the reward obtained was also negatively correlated with P-F1 (*P* < 0.01; Fig. 2b). These negative associations remained significant after controlling for age, gender, and education level of the participants (*P* < 0.01). On the other hand, the second psychiatric factor (P-F2) was not significantly correlated with task performance (*Ps* > 0.05; Fig. 2ab). Moreover, we did not find any significant associations between the reaction time in the decision-making task and the two psychiatric factors (*P* > 0.05; Fig. 2c). Given that P-F1 is mainly associated with obsessive-compulsive traits and P-F2 is mainly associated with depression and anxiety, these results suggest that overall performance of decision-making is impaired mainly by obsessive-compulsive traits rather than depression-or anxiety-related traits.

**Fig. 2:**
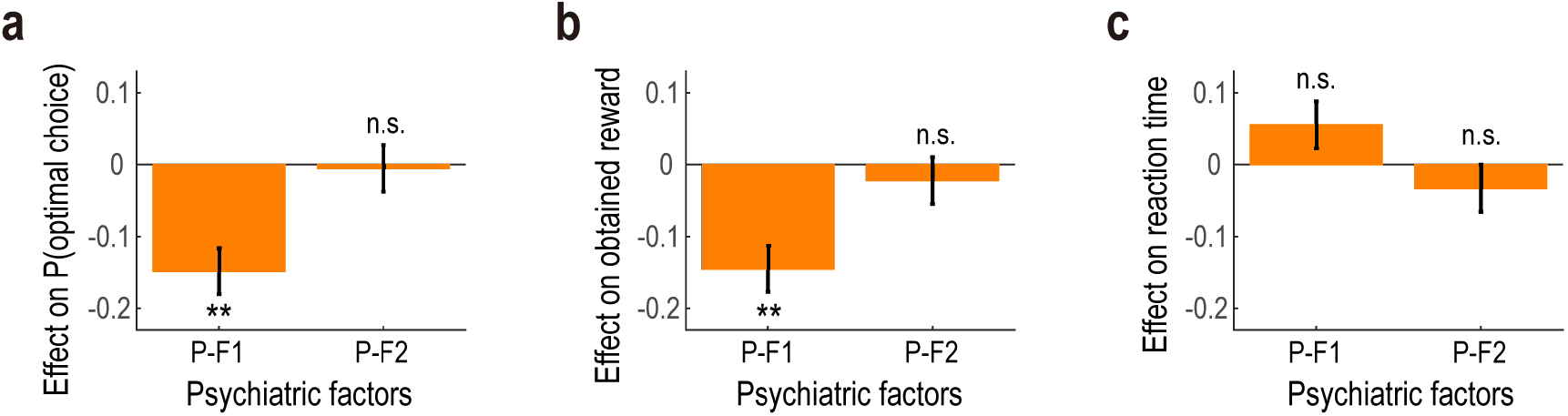
Relationship between psychiatric factors and overall task performance. (a) Effects of the psychiatric factors on the proportion of the optimal choice (mean ± SEM, *n* = 939). The mean and SEM of the effects are estimated by a linear regression analysis. \*\**P* < 0.01 (Bonferroni corrected: *α*’ = 0.01/2) and n.s., non-significant in two-tailed *t*-tests (P-F1: *β*= −0.15, *t*_936_ = −4.56, *P* < 0.001; and P-F2: *β*= −0.01, *t*_936_ = −0.16, *P* = 1.000). *P-F1*, the first psychiatric factor mainly associated with obsessive-compulsive traits; and *P-F2*, the second factor mainly associated with depressive and anxious traits. (b) Effects of the psychiatric factors on the obtained reward (*n* = 939; P-F1: *β*= −0.14, *t*_936_ = −4.46, *P* < 0.001; and P-F2: *β*= −0.02, *t*_936_ = −0.68, *P* = 0.988). The format is the same as in (a). (c) Effects of the psychiatric factors on the reaction time (*n* = 939; P-F1: *β*= 0.06, *t*_936_ = 1.69, *P* = 0.181; and P-F2: *β*= −0.03, *t*_936_ = −1.01, *P* = 0.627). The format is the same as in (a).

### Psychiatric symptoms and decision-making processes: model-neutral regression analysis

We next questioned how the first psychiatric factor (P-F1), mainly associated with obsessive-compulsive traits, is related to specific computational processes in decision-making. We assessed participants’ behavioural pattern in the decision-making task, and then tested whether and if so, how the behavioural pattern was modulated by P-F1.

The reinforcement learning account of reward-seeking decision-making predicts that an individual’s behaviour is driven by reward feedback i.e., choices that lead to reward delivery are reinforced and therefore are more likely to be selected in the next occasion. In addition, some studies have reported individuals’ behaviour is also guided by their past choices^36–38^. Importantly, the positive effects of the choice trace can be interpreted as “choice perseverance”, “choice stickiness”, or “decision inertia”. On the other hand, the negative effects of the choice trace indicate that an individual is likely to choose options that have not been chosen previously for a while. The preference for the previously unchosen options can be thought to reflect uncertainty-driven exploration, as values of the unchosen options are evidently uncertain.

To examine these effects, we employed a generalised linear mixed-effect model (GLMM) analysis. The regression model used in this study allows us to test for how participants’ behaviour was affected by past rewards and past choices (i.e., the main effects)^38–40^, and how these main effects were mediated by the psychiatric factors (i.e., the interaction effects)^26^, while controlling for age, gender and education level. We first examined the main effects of reward and choice history on the behaviour, independent of the psychiatric factors. We found a positive impact of reward history (*P* < 0.01; Fig. 3a), consistent with the prediction of reinforcement learning account. Furthermore, the effect of choice history on the behaviour was found to be significantly negative (*P* < 0.01; Fig. 3a), suggesting that participants’ choice pattern was at least in part governed by uncertain-driven exploration.

**Fig. 3:**
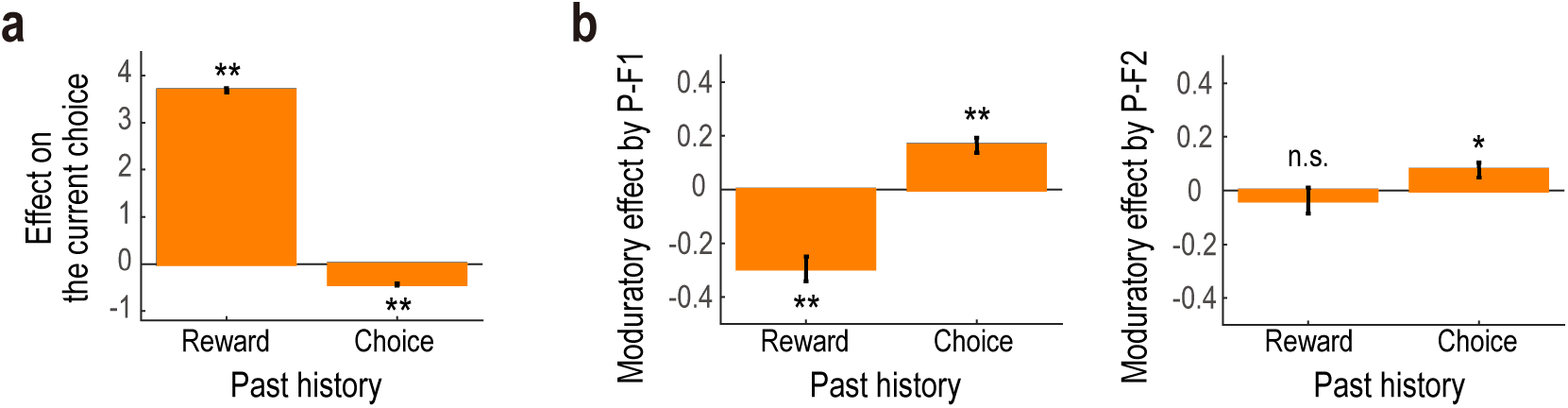
Model-neutral regression analysis about the relationship between psychiatric factors and decision-making process. (a) Effects of past rewards and past choices on the current choice independent of psychiatric factors (mean ± SEM, *n* = 461,679). The mean and SEM of the effects are estimated by a generalised linear mixed-effect model (GLMM) analysis. \*\**P* < 0.01 (Bonferroni corrected: *α*’ = 0.01/2) in two-tailed *t*-tests (Reward: *β*= 3.69, *t* _461,625_ = 80.17, *P* < 0.001; and Choice: *β*= −0.43, *t*_461,625_ = −16.50, *P* < 0.001). (b) Modulations of the reward and the choice history effects by the psychiatric factors (mean ± SEM, *n* = 461,679). The mean and SEM of the modulation effects are estimated by the same GLMM as in (a). *Left*, modulatory effects by the first psychiatric factor (*P-F1*); and *right*, modulatory effects by the second psychiatric factor (*P-F2*). \*\**P* < 0.01 (Bonferroni corrected: *α*’ = 0.01/4); and \**P* < 0.05 (Bonferroni corrected: *α*’ = 0.05/4) in two-tailed *t*-tests (Reward x P-F1: *β*= −0.29, *t* _461,625_ = −6.22, *P* < 0.001; Choice x P-F1: *β*= 0.16, *t*_461,625_ = 6.08, *P* < 0.001; Reward x P-F2: *β*= −0.04, *t* _461,625_ = −0.76, *P* = 1.000; Choice x P-F2: *β*= 0.08, *t*_461,625_ = 2.77, *P* = 0.022).

We next examined the interaction effects i.e., how the effects of past rewards and choices on the behaviour are modulated by the first psychiatric factor (P-F1), mainly associated with obsessive-compulsive traits. Both the reward and the choice of history effects were found to be attenuated by P-F1 (Fig. 3b). Specifically, the regression coefficient of the interaction term between the reward history and P-F1 was significantly negative (*P* < 0.01; Fig. 3b, *left*), indicating that the positive effect of past rewards on the behaviour was inhibited by P-F1. Moreover, the regression coefficient of the interaction term between the choice history and P-F1 was positive (*P* < 0.01; Fig. 3b, *left*), indicating that the negative effect of past choices was inhibited by the psychiatric factor. On the other hand, we found less robust interaction effects between the reward/choice history and the second psychiatric factor (P-F2) associated with depression and anxiety (Fig. 3b, *right*). These findings are consistent with the notion that the obsessive-compulsive traits diminish reward-driven behaviour and preference for previously unchosen options (i.e., the negative effect of the choice trace), which can be related to uncertainty-driven exploration in decision-making.

### Psychiatric symptoms and decision-making processes: a model-based analysis

In the previous model-neutral regression analysis, we have shown that the effects of past rewards and past choices were inhibited by the first psychiatric factors (P-F1) mainly associated with obsessive-compulsive traits. To further specify what computational processes underlie the behavioural pattern, we fitted a family of formal computational models to the participants’ behaviour and then correlated the best model’s parameters with P-F1 (see Methods and Table 1 for details about the computational models).

**Table 1.** Results of the model comparison. Results of the Bayesian model selection are described. The best-fitted model is shown in bold. See Methods and the ref^47^ for details.

|  | Model ID | # Parameter | Model Prob. | Exceedance Prob. |
| --- | --- | --- | --- | --- |
| Conventional RL models | RL1a | 2 | 0.009 | 0.000 |
|  | RL1b | 3 | 0.077 | 0.000 |
| **RL models with**<br>**choice-trace effect** | RL2a | 4 | 0.076 | 0.000 |
|  | **RL2b** | **5** | **0.642** | **1.000** |
| RL models with<br>asymmetric learning rates | RL3a | 3 | 0.006 | 0.000 |
|  | RL3b | 4 | 0.190 | 0.000 |

The first model is a conventional reinforcement learning model (RL1a), known as Q-learning. In this model, as shown in the above regression analysis, choices that lead to reward delivery are reinforced (Fig. 3a) i.e., the value of the chosen option is updated by reward prediction error and the updated value guides the individual’s next choice. Importantly, this model has two parameters. One is the learning rate which governs how much value is updated in proportion to the reward prediction error. The other is inverse-temperature which governs the sensitivity of the choice to option values (i.e., choice stochasticity). Note that the higher value of the inverse-temperature means that the individual is more likely to choose options with a higher value, while lower inverse-temperature means the individual’s decisions are more random. In other words, the parameter controls the degree of random-exploration. In addition, following previous studies^41–44^, we consider another model including forgetting of values of unchosen options (RL1b), known to provide a better account of organisms’ behaviours and tonic/phasic activity of dopamine neurons.

Given the results that participants’ behaviour is driven not only by past rewards but also by their past choices (Fig. 3a), in the second class of reinforcement learning models (RL2a and RL2b), we assumed that an individual takes into account “choice trace” in their decision-making (note: RL2a and RL2b differ in whether forgetting of unchosen value is included or not). Here, a critical parameter, called choice-trace weight, determines how much past choices affect the individual’s current behaviour. Specifically, individuals with a positive choice-trace weight are likely to repeat the choice they recently selected. On the other hand, individuals with negative choice-trace weight tend to avoid an option recently chosen, which can be interpreted as uncertainty-driven exploration. Notably, it is observed that the choice-trace effects are often confounded with the effects of differential (asymmetric) learning rates for positive and negative reward prediction errors^45,46^. We, therefore, constructed another class of models (RL3a and 3b) which introduce asymmetric learning rates instead of the choice-trace effect.

In this study, we evaluated the competing models’ goodness of fit by using Bayesian Model Selection^47^, where a higher exceedance probability closer to one indicates a better fit. Notably, before applying these models to the actual participants’ data, we confirmed, based on simulated data, that the competing models are identifiable given the current experimental settings^48^ (see “confusion matrix” in Fig. S2a). This means that each of these models captures unique behavioural patterns.

The Bayesian model selection revealed that a model including choice-trace weight (RL2b) provided a better fit compared to the alternative models (Fig. 4a and Table 1). Exceedance probability^47^ of the best model, RL2b, was almost one. To further validate the model fitting result, we generated simulation data based on the best-fitted model and the parameters^48^. In the simulated data, we confirmed that the parameter estimates can be successfully recovered by the model fitting (Fig. S2b) and that the results of the model-neutral regression analysis can be replicated (Fig. S2c), implying that the model and the parameters reliably capture meaningful computational processes^48^. In sum, consistent with the results of the regression-based analyses, the model-based analysis shows that participants’ behaviour was guided by past rewards as well as past choices.

**Fig. 4:**
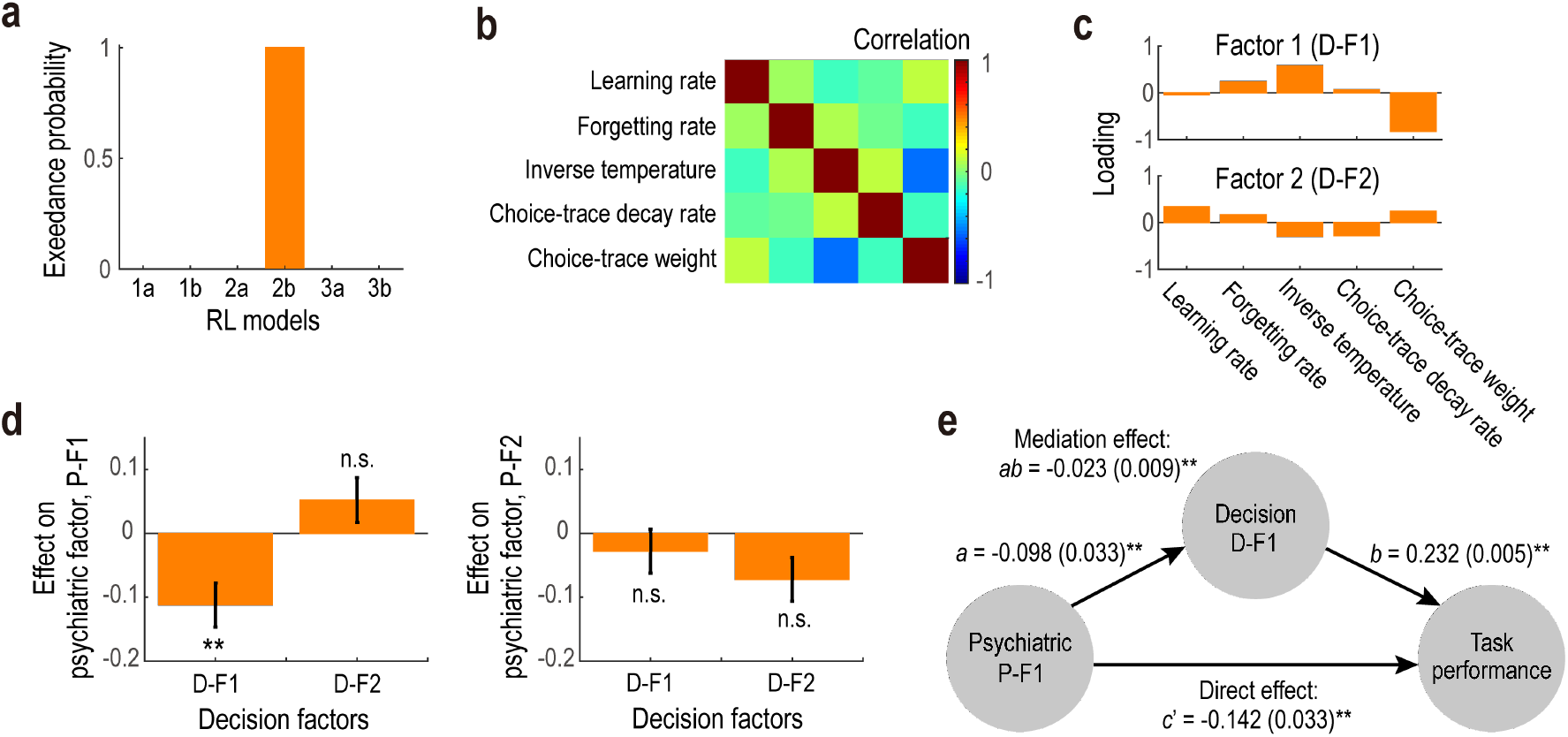
Computational model-based analysis about the relationship among psychiatric factors, decision-making process and overall task performance. (a) Results of the model comparison. Each bar denotes exceedance probability in Bayesian model selection (higher values close to *one* indicate a better fit). (b) Cross-correlation of decision parameters in the best-fitted model (i.e., RL2b). (c) Results of the factor analysis on the decision parameters. *Top*, loadings of the first decision factor (D-F1); and *bottom*, loadings of the second decision factor (D-F2). (d) Relationship between the decision factors (D-F1 and D-F2) and the psychiatric factors (P-F1 and P-F2) (mean ± SEM, *n* = 939). The mean and SEM of the effects are estimated by a linear regression analysis. \*\**P* < 0.01 (Bonferroni corrected: *α*’ = 0.01/4) and n.s., non-significant. *Left*, two-tailed *t*-tests (D-F1: *β*= −0.11, *t*_932_ = −3.22, *P* = 0.005; and D-F2: *β*= 0.05, *t*_932_ = 1.49, *P* = 0.544). *Right*, two-tailed *t*-tests (D-F1: *β*= −0.03, *t*_932_ = −0.81, *P* = 1.000; and D-F2: *β*= −0.07, *t*_932_ = −2.09, *P* = 0.146). (e) Mediation path diagram for the first psychiatric factor (P-F1), the first decision factor (D-F1) and the task performance (*n* = 939). The mean path coefficients (i.e., *β* values) are shown with SEM (in parentheses). Here, task performance is defined as the proportion of the optimal choice. The path *a* denotes influence of P-F1 on D-F1 (*P* = 0.005, *Bootstrap* test), while the path *b* depicts the influence of D-F1 on the task performance (*P* = 0.003, *Bootstrap* test). Importantly, the path *ab* indicates whether the coupling between P-F1 and the task performance was mediated by D-F1 (*P* = 0.006, *Bootstrap* test). The path *c* reflects the direct relationship between P-F1 and the task performance (*P* < 0.001, *Bootstrap* test), controlling for the mediation effect of D-F1. Please see the ref^49,50^ for details. \*\**P* < 0.01.

We then aimed to relate the model’s decision parameters to the psychiatric factors. We first extracted the decision-related dimensions underlying the five parameters in the best-fitted model (Fig. 4b). A factor analysis revealed two decision factors. The first one (D-F1) is characterised by a higher negative loading of choice-trace weight and positive loading of inverse-temperature (Fig. 4c, *top*). The second one (D-F2) is characterised by moderate positive loading of learning rate (Fig. 4c, *bottom*). By regressing the two decision factors (D-F1 and D-F2) against the first psychiatric factor (P-F1) mainly associated with obsessive-compulsive traits, we found that D-F1 had a significantly negative association with P-F1 (*P* < 0.01; Fig. 4d, *left*), while D-F2 did not (*P* > 0.05; Fig. 4d, *left*). Notably, in this regression analysis, potentially confounding effects of age, gender, education level and the other psychiatric factor (P-F2) were controlled. On the other hand, neither D-F1 nor D-F2 was significantly correlated with the P-F2 (*Ps* > 0.05; Fig. 4d, *right*). Given that the higher inverse-temperature corresponds to a lower choice stochasticity (i.e., a lower degree of random exploration) and that negatively higher choice-trace weight can reflect a higher degree of uncertainty-driven exploration, the negative relationship between D-F1 and P-F1 implies that higher obsessive-compulsive traits are associated with particular decision-making strategies i.e., a higher degree of random-exploration and a lower degree of uncertainty-driven exploration.

### Relationship among psychiatric symptoms, decision-making strategies and task performance: a mediation analysis

So far, we have shown that the overall task performance was negatively coupled with the psychiatric factor (P-F1) mainly associated with obsessive-compulsive traits and that P-F1 was negatively correlated with the decision factor (D-F1) that governs choice stochasticity and choice-trace weight (possibly reflecting random and uncertainty-driven exploration strategies respectively). We further tested whether the negative coupling between P-F1 and the task performance was mediated by D-F1. A formal mediation analysis^49,50^ revealed a significant mediation effect of D-F1 (*P* < 0.01; Fig. 4e), after controlling for effects of age, gender, education level, the other psychiatric factor (P-F2), and the other decision-factor (D-F2). These findings together are consistent with the idea that obsessive-compulsive traits impair the overall performance of decision-making through the change of computational processes that can govern random and uncertainty-driven exploration strategies.

Finally, in order to clarify the role of the psychiatric factor (P-F1) beyond the mere total score of Obsessive-Compulsive Inventory (OCI)^51^, we conducted the above mediation analysis with the total score of OCI instead of P-F1. Although coupling between the OCI score and the task performance remained significant (*P* < 0.01; Fig. S3b), we found no significant evidence for the association between the OCI score and the decision factor (D-F1) (*P* > 0.05; Fig. S3b). Moreover, the mediation effect of D-F1 on the coupling between the OCI score and the task performance was found to be insignificant (*P* > 0.05; Fig. S3b). These findings could imply the importance of the trans-diagnostic approach in computational psychiatry^26^.

## Discussion

The present study provides a mechanistic account of how psychiatric symptoms are related to impairments in value-based decision-making, by using a large-scale online behavioural experiment with healthy participants. Combining computational modelling with factor and mediation analyses, we found that there are at least two dimensions underlying multiple psychiatric symptoms and that one dimension, mainly associated with obsessive-compulsive traits, is negatively correlated with the decision-making task performance. We also found that the negative correlation is mediated by individual differences in choice stochasticity (i.e., random exploration) and propensity to choose options previously unchosen, which may reflect uncertainty-driven exploration. The present findings go beyond the results from other computational psychiatry studies in that we revealed the tripartite relationship among a particular dimension of psychiatric symptoms, specific computational processes in value-based decision-making, and overall task performance.

Based on the large-scale online questionnaire data collected from healthy participants, we have identified at least two dimensions underlying psychiatric symptoms. The first one is mainly associated with obsessive-compulsive traits, while the second one is mainly associated with depressive and anxious traits. This finding was replicated in our independent sample and was broadly consistent with a large-scale longitudinal face-to-face survey on mental disorders involving healthy and patient populations^31^. The face-to-face survey study suggests that common mental disorder in adults can be characterised by three factors: externalising disorder (expressing alcohol, cannabis, and drug disorder), internalising disorder (expressing major depression and anxiety disorder) and thought disorder (expressing obsessive-compulsive disorder and schizophrenia). The first and second psychiatric dimensions, that we identified, would correspond to thought disorder and internalising disorder, respectively. Our findings together with the previous survey result have implications in studies of obsessive-compulsive disorder. One of the important open questions in this field is to what extent obsessive-compulsive disorder is caused by elevated anxiety^52^. Some researchers believe compulsive behaviour is elicited by the motivation to reduce anxiety, while obsessive-compulsive and anxiety-related disorders are classified in different categories in DSM5 (5th edition of Diagnostic and Statistical Manual of Mental Disorders)^53^. Our finding is consistent with the possibility that obsessive-compulsive disorder is basically a dissociable psychological process from anxiety, although further studies are required in order to collect more promising evidence.

More broadly, the replicability of our questionnaire data and its consistency with the face-to-face survey involving the patient population validates the use of online experimentation with healthy participants. In mental disorders comorbidity rate is known to be high^31,32^, and correspondence between categorical diagnosis of mental disorder and psychological processes seems to be not one-to-one i.e., multifinality and equifinality^33,34^. The high comorbidity, multifinality and equifinality make it difficult to examine the relationship between particular psychiatric symptoms and impairment of decision-making, thereby requiring careful control for confounding effects of the other symptoms. Of note, control for the confounds is virtually difficult in typical studies with a small sample size, where tens of patients and healthy participants are recruited for the comparison of decision-making performance. The rigorous comparison between participants usually requires hundreds of participants with various psychiatric symptoms. To overcome the difficulty, online experiments with healthy participants is emerging as a popular tool in psychology^54^ and psychiatry^35^. The present study supports the utility of online experiments for the investigation of psychological mechanisms underlying mental disorders.

Comparison between the two psychiatric dimensions and overall performance of the decision-making task revealed that the first dimension (with high loading from obsessive-compulsive traits), but not the second dimension (with high loading from depressive and anxious traits), is negatively correlated with the task performance. It is worth noting that the first psychiatric dimension contains components not only from obsessive-compulsive traits but also from other psychiatric symptoms. The dimension is therefore not obsessive-compulsive traits itself, but one-dimension consisting of various psychiatric symptoms. Previous studies have reported that impairment of decision-making was accompanied by various psychiatric symptoms such as obsessive-compulsive disorder^20,21^, depression^12^, anxiety^32^, and schizophrenia^26^. One interpretation of these mixed reports is that all the symptoms are most likely related to decision-making. The other possible interpretation is that there are hidden dimensions underlying the multiple symptoms and that only one of them is essentially coupled with impairments of decision-making. It was, however, difficult to tease out the two interpretations, given the high comorbidity of psychiatric symptoms, the small sample size, and the lack of control analysis for confound effects of other symptoms. The present study may support the latter interpretation, based on the large-scale online experiment with decision-making task and multiple questionnaires about many psychiatric symptoms, i.e., one dimension of the psychiatric symptoms consisting mainly of obsessive-compulsive traits are coupled with impairments of decision-making.

We also demonstrated that the first psychiatric dimension (mainly associated with obsessive-compulsive traits) is negatively associated with the lower degree of choice stochasticity and the higher propensity to choose options previously unchosen. The choice stochasticity is often related to a random exploration^9,11^. Moreover, the choice preference for recently unchosen options can be thought to reflect uncertainty-driven exploration, as options that have not been chosen previously for a while are evidently uncertain about their values^9^. Taken together, our findings are consistent with the view that psychiatric symptoms mainly associated with obsessive-compulsive traits are linked to different types of exploration (i. e. a higher degree of random exploration and a lower degree of uncertainty-driven exploration).

We believe our findings could have implications for understanding the psychological mechanisms underlying the obsessive-compulsive disorder. Core symptoms of the obsessive-compulsive disorder are generally characterised by compulsive rituals such as repetitive adverse (e.g., excessive hand-washing) and obsessive checking behaviours (e.g., endless rechecking if the door is locked). Our findings of the association between obsessive-compulsive traits and the decrease in the propensity to choose options previously unchosen may suggest a novel and parsimonious psychological mechanism underlying the obsessive-compulsive disorder. That is, whereas typical people tend to choose an option that has not been chosen previously in order to resolve the uncertainty (i.e., uncertainty-driven exploration), people with higher obsessive-compulsive traits tend to keep choosing the same option even though its value is no longer uncertain. In other words, failure to utilise the information about uncertainty or decline of the motivation to reduce uncertainty could be a promising and parsimonious account for the core symptoms in obsessive-compulsive disorder. Interestingly, this account is broadly consistent with the recent study^55^, suggesting that patients with obsessive-compulsive disorder correctly estimate confidence or uncertainty of their belief about the environmental state but fail to utilise the knowledge to appropriately update their belief. This account is also broadly coherent with the long-lasting hypothesis about the endophenotype of obsessive-compulsive disorder that patients of the disorder lose the cognitive flexibility (e.g., response inhibition) observed in classical experimental tasks such as Reversal Learning Task and Go/NoGo Task^52,56,57^. Furthermore, it is worth noting that our proposed account is not mutually exclusive with another influential account of obsessive-compulsive disorder where the contribution of a flexible model-based reinforcement learning to the decision-making is reduced^25,26^. An interesting avenue for future research would be to uncover a relationship between declines of uncertainty-driven behaviours and model-based reinforcement learning with respect to obsessive-compulsive traits.

Exploration strategies have been demonstrated to be linked not only to obsessive-compulsive disorder but also with other mental disorders. For example, one study reported that uncertainty-driven exploration is reduced in patients with schizophrenia^29^. Furthermore, in healthy participants, depressive traits have been found to correlate with the degree of random exploration^28^. A possible account that can unify these divergent findings would be that the exploration strategies are related to one dimension of psychiatric symptoms that we found in the present study. This was associated mainly with obsessive-compulsive traits as well as with depressive and schizotypal traits to some degree. Given the sparsity of studies on this issue, more evidence is needed for a comprehensive understanding of the relationship between exploration strategies and mental disorders.

Finally, a formal mediation analysis revealed that the negative coupling between one dimension of psychiatric symptoms (mainly associated with obsessive-compulsive traits) and overall performance of decision-making is mediated by a computational process that can govern choice stochasticity and preference for previously unchosen options, which can be related to random and uncertainty-driven exploration strategies, respectively. Various psychiatric symptoms are known to be accompanied by impairments in the overall performance of decision-making, and recent studies in computational psychiatry have begun to elucidate a correspondence between psychiatric symptoms and specific computational processes in decision-making^20,21^. However, surprisingly few studies to date have examined the tripartite relationship among particular psychiatric symptoms, the overall performance of decision-making, and specific computational processes carried out in the brain. The present study goes beyond the previous computational psychiatry studies by providing an overview of the tripartite relationship.

One caveat of this study would be that the conclusion could depend on the decision-making task used. For example, exploration strategies could not play a pivotal role in stable environments such as multi-armed bandit task with constant reward probabilities. Furthermore, in environments with changing volatility, different computations underlying decision-making and learning might play an essential role^58,59^. Another caveat is that, while we relate choice stochasticity and preference for previously unchosen options to random and uncertainty-driven exploration strategies respectively, more sophisticated experimental tasks are required for in-depth examinations of the nature of the exploration strategies^8,10,12,60^. Furthermore, while we have shown the correlation between one dimension of psychiatric symptoms and particular computational processes in decision-making, we have left open the question as to how these two are causally related. That is, psychiatric symptoms can cause changes in computational processes or vice versa. An interesting future direction would be to conduct intervention studies in order to further examine the causal relationship.

To conclude, by combining a large-scale online experiment with model-based data analysis, we propose a mechanistic account of how negative coupling between psychiatric symptoms and decision-making performance is mediated by hidden computational processes carried out in our brain. Computational processes that govern exploration strategies may mediate the negative coupling. By untangling interdependent relationships among psychiatric symptoms, psychological/computational processes and overall decision-making performance, the present study provides an important scaffold for a comprehensive understanding of computational principles of information processing in patients with mental a disorder.

## Methods

The study was approved by the ethics committee in the Department of Psychology, Graduate School of Informatics, Nagoya University (ID: NUPSY-171027-K-01).

### Participants

1200 participants (600 females; age range, 20-69 years; age mean ± SD, 48.10 ± 11.58) completed the online experiment consisting of a reward-seeking decision-making task and questionnaires. They gave a response on more than 90% of trials in the decision-making task and provided answers to all the questionnaires. All the data were collected using an online research company, Cross Marketing Inc. (http://global.cross-m.co.jp/). Note that the research company did not play any roles in designing the experiment, analysing the data or writing of the manuscript. All the participants were native Japanese speakers and pre-assessed to exclude those with any previous history of neurological/psychiatric illness. All participants gave their informed consent online by clicking ‘I Agree’ after reading instructions of the experiment, and received a monetary reward depending on their performance in the decision-making task in addition to the participation fee of 500 Japanese *yen* (see Reward Payment for details). We did not use any statistical methods to predetermine the sample size, and our sample size selection was motivated by those used in the previous studies^26^.

### Exclusion Criteria of the Participants

In order to ensure the data quality in the online experiment, following the previous studies^26,61^, we excluded 261 participants in total by careful assessments. First, we excluded 223 participants who were not serious about the questionnaire. To find out such participants, we included a catch item, “If you have carefully read the questions so far, please select ‘a little’ as your answer”, in the questionnaire (see Experiment for details). If participants failed to choose the appropriate response, they were excluded from the subsequent analysis. Two participants were excluded as they did not provide information about their education level. Finally, we excluded 36 participants who chose one option on more than two-thirds of the trials in the decision-making task. Choices of one option over the course of trials (e.g., more than *two-thirds* of the trials) are highly irrelevant in terms of the task demands, given that in this task the optimal agent would choose all the three options with almost the same frequencies (Fig. 1b, *right*). In other words, frequencies of trials in which each of the three options had the highest reward probability were almost the same in the task (Fig. 1b, *right*). We, therefore, used the data from the remaining 939 participants (494 females; age range, 20-69 years; age mean ± SD, 47.90 ± 11.70) in the subsequent data analyses.

### Experiment

In our online experiment, participants performed a reward-seeking decision-making task (a conventional three-armed bandit task) and then answered questionnaires about psychiatric symptoms as well as their socio-economic status. Prior to the main test session, participants completed a practice session of the decision-making task (50 trials).

In the decision-making task, participants chose among the three options repeatedly (Fig. 1a), corresponding to 500 trials (they had a one-minute break every 100 trials). In each of the 500 trials, they received a reward (100 Japanese *yen* or 0 *yen*) depending on the probability assigned to the chosen option. As the reward probability for each option was unknown to the participants and changed dynamically across trials (Fig. 1b), they were required to keep track of the probabilities over the course of the experimental task to maximise their reward earnings. Prior to the experiment, they were told that “the reward probabilities may change during the task”. Notably, all the participants were confronted with the same reward probabilities, so that we could exclude the possibility that differences in the changing pattern of reward probabilities can account for any individual differences in the participants’ behaviour.

At the beginning of each trial, participants were asked to make a choice among the three options by clicking one of the fractal stimuli within 6 s (Decision phase; Fig. 1a). In this task, the three options (*green*, *red* and *blue*) were positioned to the left, middle and right of the screen respectively in every trial. After making a response, the chosen option was highlighted by a yellow frame (Confirmation phase, 1 s), and then an outcome of the choice (100 yen or 0 yen) was revealed to the participants (Feedback phase, 1.5 s). If no response was made in the decision phase, the remaining phases were skipped, and the participants moved to the next trial. During this task, participants failed to make a response only in 0.87 ± 1.75% of trials (mean ± SD).

After the decision-making task, participants were asked to answer the Japanese versions of the following questionnaires: Schizotypal Personality Questionnaire Brief^62,63^, Obsessive-Compulsive Inventory^51^, Self-Rating Depression Scale^64^, State-Trait Anxiety Inventory^65^, Barratt Impulsivity Scale^66^, and Socioeconomic status^67^. We presented these questionnaires in the order above to all the participants, to ensure that any individual differences in the self-reported psychiatric symptoms cannot be attributed to any potential order-effects. Notably, to identify participants who were not serious about the questionnaires, we added one catch question, “If you have carefully read the questions so far, please select ‘a little’ as your answer”, to Obsessive-Compulsive Inventory (see also Exclusion Criteria of the Participants).

### Reward Payment

In addition to a participation fee of 500 Japanese *yen*, participants obtained an extra reward depending on their performance in the decision-making task. Amount of the performance-based reward was determined as follows: at the end of the experiment, the computer randomly selected one trial, and the outcome of the trial was implemented. That is if the participant received a reward in that trial they obtained 100 *yen*. Importantly, since participants did not know which a trial would be selected for the monetary reward, they should have treated every trial as if it were actually being implemented. The reward they earned was paid by a coupon which could be used in popular Japanese online stores including Amazon Japan (https://www.amazon.co.jp/) and Rakuten Ichiba (https://www.rakuten.co.jp/).

### Data Analysis

The data were basically analysed by using MATLAB R2018b on iMac (Retina 5K, 27-inch, 2019; Mac OS X 10.14.4).

#### Factor analysis on the questionnaire data

We conducted a factor analysis on the participants’ responses to 160 individual items in the six questionnaires. As responses to the individual items are categorical, the factor analysis was performed on the Polychoric correlation matrix (Fig. 1d). The number of factors, two, was determined by a prior hypothesis based on the large-scale longitudinal face-to-face survey on healthy and patient populations^31^. This large-scale face-to-face survey^31^ suggests that common mental disorders in adults can be characterised by three factors: externalising disorder (expressing alcohol, cannabis, and drug disorder), internalising disorder (expressing major depression and anxiety disorder) and thought disorder (expressing obsessive-compulsive disorder and schizophrenia). Given that psychiatric symptoms measured in our questionnaires (e.g., schizotypal traits, obsessive-compulsive traits, anxious traits, and depressive traits) would perhaps correspond to the two factors in the survey (*thought disorder* and *internalising disorder*), we hypothesised that there would exist two factors underlying our questionnaire data. Consistent with this hypothesis, the scree-plot derived from the Polychoric correlation matrix indicated that slopes of the eigen-values were less steep at the 3^rd^ and latter factors (Fig. S1b), implying that retainment of additional factors yields little benefit. The resulting pattern of the factor loadings (Fig. 1e) is found to be consistent with the large-scale face-to-face survey^31^. Moreover, it is worth noting that the pattern of the factor loadings does not change even if we assume there exist three (rather two) factors underlying the symptoms (Fig. S1c). Given the number of factors, the factor loadings were estimated by the Maximum likelihood method, and Varimax rotation was applied. We then estimated factor scores for each participant by using Bartlett algorithm. In this study, we employed an orthogonal rotation method, Varimax rotation, as we aimed to serve the estimated factor scores as mutually dissociable exploratory variables in linear regression analyses. As polychoric correlation is not implemented in MATLAB, the analyses were performed by an *R* function (version 3.6.0), *fa*.

#### Psychiatric symptoms and overall task performance: linear regression analyses

In order to examine the relationship between psychiatric symptoms and overall task performance, we conducted linear regression analyses. In other words, we regressed the two psychiatric factor scores (*F1* and *F2*), identified by the factor analysis, against the task performance (*P*): *P* ~ 1 + *F1* + *F*2, where the regression model is specified by Wilkinson notation (mainly used in R and MATLAB). Statistical significance of the regression coefficients was tested with Bonferroni multiple-comparison correction by the number of explanatory variables of interest (i.e., two: *F1* and *F2*). In the first analysis, the task performance was defined as the participant’s proportion of the optimal choice (i.e., choice of the option that had the highest reward probability), while in the second analysis the performance was defined as the proportion of trials rewarded. Furthermore, in the additional analyses, we incorporated the participants’ age, gender (coded as male: 1; female: 2) and education-level (coded as a junior high school diploma: 1; high school diploma: 2; technical school diploma: 3; vocational school diploma: 4; associate degree (community college diploma): 5; bachelor’s degree: 6; and a master’s or doctorate degree: 7) as variables of no-interest. Note that all the variables were *z*-normalised and fed into the regression analyses.

#### Psychiatric symptoms and decision-making processes: model-neutral regression analysis

In order to examine how past rewards and choices affect the participant’s current behaviour and how these effects are modulated by the psychiatric factors, we conducted a Generalised Linear Mixed Model (GLMM) analysis by using a MATLAB R2018b function, *fitglme*, with the restricted maximum pseudo-likelihood estimation. Following the previous studies using three-armed bandit task^39,68^, we performed three separate logistic regression models, one for each choice option (*X*, *Y* and *Z*).

For one option *X*, the GLMM was defined as follows (in the Wilkinson notation):

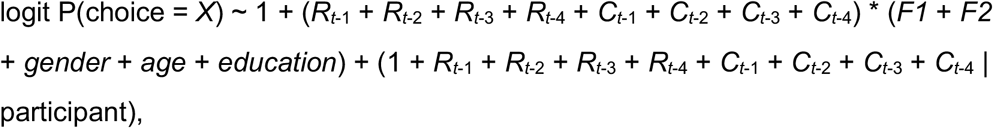

where *R*_*t*−*τ*_ and *C*_*t*−*τ*_ denote recent past rewards and recent past choices respectively (trials *t*−1, *t*−2, *t*−3, and *t*−4), and *F1* and *F2* represent the two psychiatric factors. As in the previous studies^39,68^, *R*_*t*−*τ*_ was coded as 1 if the participant chose *X* and obtained a reward on trial *t* − *τ*, −1 if they chose *Y* or *Z* and obtained a reward, and 0 if there was no reward. *C*_*t*−*τ*_ was coded as 1 if the participant chose *X* on trial *t* − *τ*, and −1 if they chose *Y* or *Z.* In this model, the participants’ demographic information (i.e., z-normalised gender, age and education-level) was controlled, and the term (. | participants) indicates within-participant variables taken as random-effects (i.e., allowed to vary between participants). The model provided us with a set of regression coefficients and the covariances. Here, we were interested in the fixed-effects of past rewards and past choices and their interactions with the psychiatric factors. The total effect of past rewards over the past *four* trials can be derived by 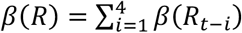, where *β*(*R*_*t*−*i*_) denotes the regression coefficient of the variable *R*_*t*−*i*_; and the variance of the total effect can be computed by 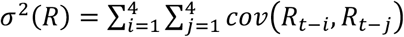 where *cov*(*R*_*t*−*i*_, *R*_*t*−*j*_) denotes the covariance of the regression coefficients, *β*(*R*_*t*−*i*_) and *β*(*R*_*t*−*j*_). The total effect of past choices and its variance, as well as the total interaction effects and their variances, can be derived in the same way.

The GLMMs for the other two options, *Y* and *Z*, were defined in the same manner, providing the total effects of past rewards, past choices and their interactions with the psychiatric factors. The mean effect of past rewards over the three models can be then derived by the variance-weighted mean^69^: 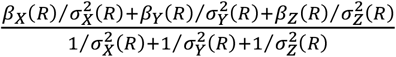, where subscripts denote each of the three models. The variance of the mean is given by 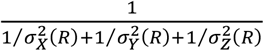 (see the ref^69^). Based on the mean effect and its variance (standard deviation), we tested the statistical significance of the effect by using two-tailed *t*-test with Bonferroni multiple-comparison correction (see the legends of Fig. 3). The same procedure was applied to the statistical tests for the effect of past choices and the effects of the interactions with the psychiatric factors.

#### Psychiatric symptoms and decision-making processes: a model-based analysis

To quantitatively capture the computational process underlying decision-making and its relationship with psychiatric symptoms, we constructed computational models and fitted them to the participants’ choice behaviours in the decision-making task (see Table 1).

##### RL1a

The first model is a conventional reinforcement learning model, called Q-learning^7^. In this model, on each trial, an agent makes a choice depending on the value of each option. That is, choice probability of an option *X* is given by

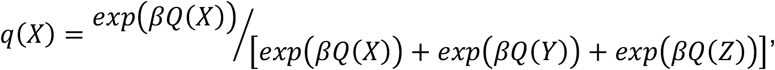

where *Q* denotes option values and the parameter, *β* ∈ [0, ∞), governs the degree of stochasticity in the choices (called inverse-temperature)^7^. Once a choice is made and the reward outcome is revealed, they update the value of the chosen option (i.e., learning).Suppose an option *X* is chosen, and then the value of the option *X* is updated as follows:

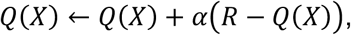

where *R* denotes reward outcome (coded 1 for the reward, and 0 for no reward) and the parameter, *α* ∈ [0,1], is the learning rate^7^. In the initial trial of the experiment, options values are set at 0.5 (as the agent seems to have no knowledge about the reward probabilities).

The other models below are variants of the conventional reinforcement learning model (i.e., RL1a). That is, the value of each option is learned through experience and governs the decision-making.

##### RL1b

This model is almost the same as RL1a but includes “forgetting”. That is, values of the unchosen options are forgotten (i.e., decayed with time)^41–44^. In other words, on each trial, an agent updates not only the value of the chosen option but also values of the unchosen options. Specifically, values of the unchosen options, say *Y* and *Z*, are updated as follows:

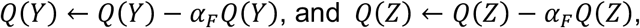

where *α*_*F*_ ∈ [0,1] denotes the forgetting rate. Note that the value of the chosen option is also updated as in RL1a.

##### RL2a

In this model, an agent takes into account her own choice-trace in the decision-making. Choice-trace of each option, which quantifies how often the option was chosen recently, is updated by the following rule:

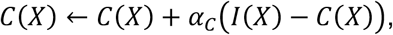

where *I*(·) is 1 if the option is chosen and 0 otherwise, and the *α*_*C*_ ∈ [0,1] is the choice-trace decay rate. Note that the choice-traces of the other two options are updated by the same rule and that the initial choice-traces are set at *zero*. The choice-traces work on the decision-making as follows:

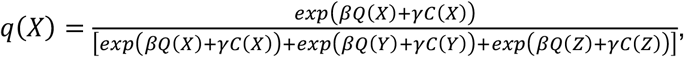

where *γ* ∈ (−∞, ∞) denotes the weight of the choice-traces. That is, the agent considers not only reward values but also choice-traces when making a decision. Importantly, individuals with positive choice-trace weights are likely to repeat the choice recently selected. On the other hand, individuals with negative choice-trace weights tend to avoid an option recently chosen.

##### RL2b

This model is almost the same as RL2a, but includes “value-forgetting”. That is, values of the unchosen options are forgotten.

##### RL3a

This model has differential learning rates for rewarding and non-rewarding outcomes. Namely, the value of the chosen option is updated as follows:

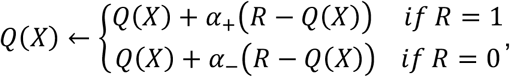

where *α*_+_ ∈ [0,1] and *α*_−_ ∈ [0,1] depict the learning rates for rewarding and non-rewarding outcomes respectively. The process of decision-making is identical to that of RL1a.

##### RL3b

This model is almost the same as RL3a but includes value-forgetting (i.e., values of the unchosen options are decayed).

##### Model fitting and comparison

To fit the computational models to each participant’s choice data, we employed a maximum a posteriori (MAP) approach incorporating prior belief about the parameter values (rather than a maximum likelihood approach in which variances of the parameter estimates are known to be inflated)^70^. Following the previous study^71^, we set prior distributions of the value learning rates, *α*, *α*_+_ and *α*_−_ at *Beta*(2,2) and the inverse-temperature, *β*, at *Gamma*(2,3). For the forgetting rate and the choice-trace decay rate, we used flatter prior distributions, *Beta*(1.2,1.2), as we did not have strong hypotheses about the parameter values a priori. A prior distribution of the choice-trace weight is set at *Norm*(0,4). For each model and each participant, we obtained a MAP estimate by the MATLAB function, *fmincon*, and then computed log model evidence by using Laplace approximations^70^. Here, note that log model evidence implicitly penalises models with more free parameters compared with those with fewer free parameters. Each model’s log model evidence was finally fed into Bayesian Model Selection^47^ for comparison of competing models’ goodness of fit.

##### Confusion matrix of model fitting

Before fitting these models to the actual participants’ data, we checked identifiability of the models given the current experimental settings^48^. That is, we tested whether each of the models captures unique behavioural patterns. To this end, we constructed a “confusion matrix”^48^ based on simulated data. If the models are perfectly identifiable, simulation data generated by one model should be best explained by the same model rather than the other models, and thus the confusion matrix should be the identity matrix. In this analysis, we first generated simulation data of 939 agents for each model. The number of agents is the same as that of the actual participants (see *Exclusion Criteria of the Participants*), and decision parameters of each agent were sampled from the prior distributions (see *Model fitting and comparison*). We then formed a confusion matrix based on exceedance probabilities of Bayesian Model Selection^47^. As a result, we found that exceedance probabilities in the diagonal cells are almost *one* (Fig. S2a), suggesting that these models are identifiable given the current experimental settings^48^.

##### Parameter recovery analysis of model fitting

After having obtained the model-fitting result, for further validation, we generated a simulation data of 939 agents whose decision parameters were extracted by the best-fitted model and parameters^48^. On the simulated data, we aimed to confirm that the parameter values can be recovered by the model fitting and that the results of the model-neutral regression analysis can be replicated. In the analyses, the parameter values were successfully recovered (Fig. S2b) and the results of the regression-based analyses on the simulation data were highly consistent with those on the actual data (Fig. S2c), implying that the model and the parameters reliably captured meaningful computational processes and their individual differences^48^.

##### Factor analysis on the decision parameters

We conducted a factor analysis of the *five* parameter estimates in the best-fitted model (RL2b). The factor analysis was performed by a MATLAB function, *factoran*. In the analysis, values of the *inverse-temperature* parameter were log-transformed due to the severe non-normality (skewness > 2 and kurtosis > 7), as suggested by the previous methodological papers on factor analysis^72,73^. Unlike the questionnaire data, the parameter estimates are continuous variables, therefore, we used Pearson correlation to form the correlation matrix. As we did not have a prior hypothesis about the number of factors underlying the decision parameters, the number was determined by Parallel analysis. We compared the scree-plot derived from the actual data with that derived from randomly generated data. The Parallel analysis indicates that eigen-values of the first and second factors in the actual data were greater than those in the random data, suggesting that there exist two underlying factors (Fig. S3a). As in the factor analysis on the questionnaire data, given the number of factors (i.e., *two*), the factor loadings were estimated by Maximum likelihood method, Varimax rotation was applied, and then factor scores for each participant were estimated using Bartlett algorithm. Notably, as mentioned in the factor analysis on the questionnaire data, we employed an orthogonal rotation method, because we intended to serve the factor scores as mutually dissociable exploratory variables in linear regression analyses.

##### Relationship between decision-related factors and psychiatric factors

We performed linear regression analyses, to examine the relationship between the two decision factors (say *D-F1* and *D-F2*) and the two psychiatric factors (*P-F1* and *P-F2*). In the first analysis, we regressed *D-F1* and *D-F2* against *P-F1*: *P-F1* ~ 1 + *D-F1* + *D-F2*. On the other hand, in the second analysis, we tested for the regression model: *P-F2* ~ 1 + *D-F1* + *D-F2*. In these models, participants’ age, gender, education level and the other psychiatric factor (*P-F1* or *P-F2*) were included as variables of no-interest, and all the variables were *z*-normalised. Finally, the statistical significance of the regression coefficients was tested with Bonferroni multiple-comparison correction by the number of explanatory variables of interest (see the legends of Fig. 4d).

#### Relationship among psychiatric symptoms, decision-making strategies and task performance: a mediation analysis

We conducted a mediation analysis to test whether the first decision factor (D-F1) mediates the negative coupling between the first psychiatric factor (P-F1) and the overall performance of decision-making task (proportion of the optimal choice). The analysis was performed by the Mediation and Moderation Toolbox^49,50^, in which the significance of the mediation effect was tested by Bootstrap procedure with 10,000 samples and all the variables were *z*-normalised. Here in this analysis, we included participants’ age, gender, education level, the second psychiatric factor (P-F2) and the second decision factor (D-F2) as variables of no-interest.

## Acknowledgements

This work was supported by the JSPS KAKENHI Grants: JP17H05933 and JP17H06022 (S.S.), JP17H06039 and JP19H04998 (Y.Y.), and 17H05946 and 18KT0021 (K.K.); and by the JST CREST Grant JPMJCR16E2 (Y.Y.).

---

**Fig. S1:**
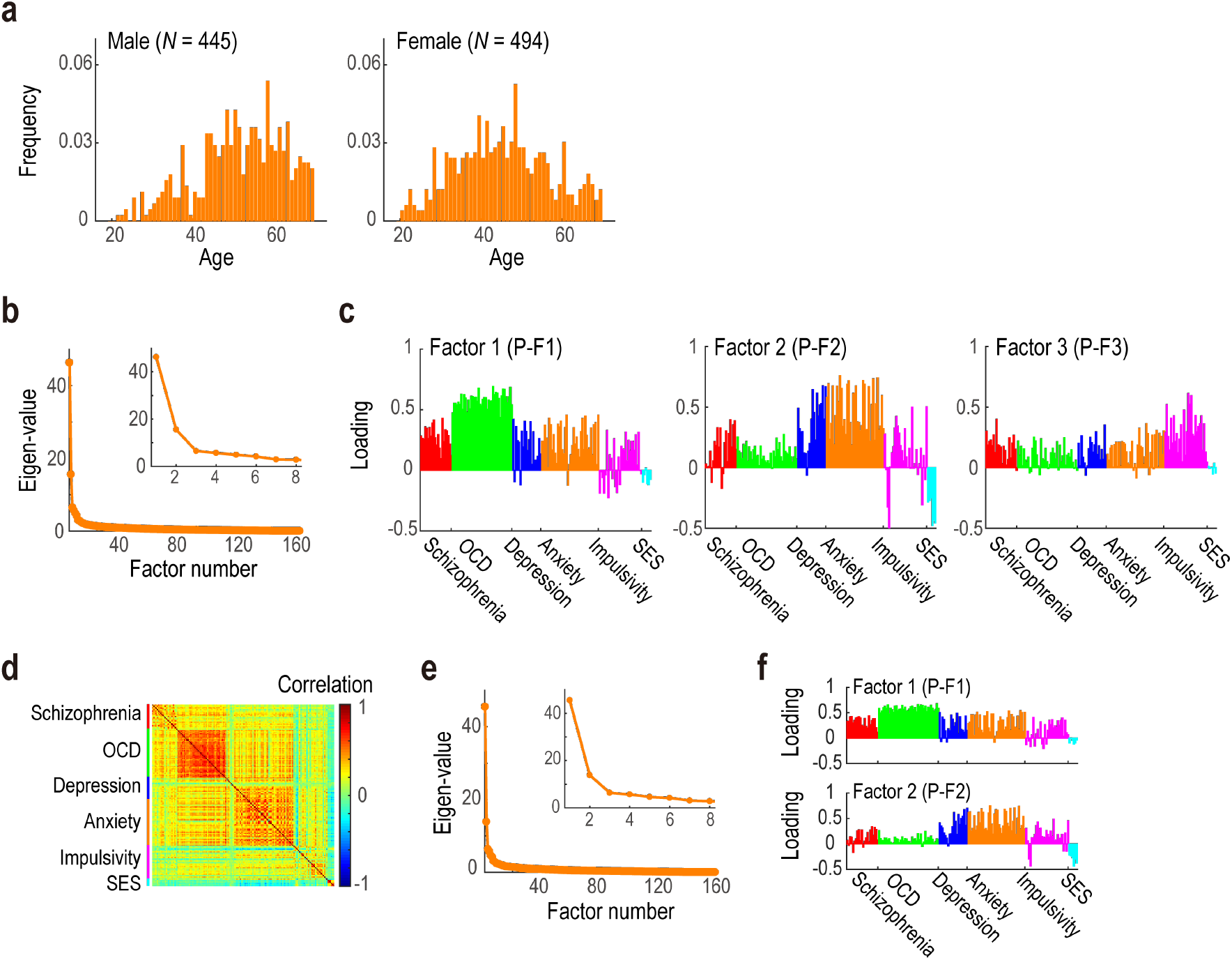
Supplementary information about the experimental settings and the questionnaire data. (a) Histograms of the participants’ age. *Left*, males (*N* = 445); and *right*, female (*N* = 494). (b) Scree-plot of the questionnaire data. We plot eigen values derived from the cross-correlation matrix (Fig. 1c). The inset shows the same data with the focus on factor 1 to 8. (c) Loadings of the questionnaire items given the assumption that three factors underlie the psychiatric symptoms. *Left*, loadings in the first factor (P-F1); *middle*, loadings in the second factor (P-F2); and *right*, loadings in the third factor (P-F3). (d) Replication of the factor analysis in an independent sample: cross-correlation matrix. The format is the same as in Fig. 1d. (e) Replication of the factor analysis in an independent sample: scree-plot. The format is the same as in Fig. S1b. (f) Replication of the factor analysis in an independent sample: loadings of the questionnaire items. The format is the same as in Fig. 1e.

**Fig. S2:**
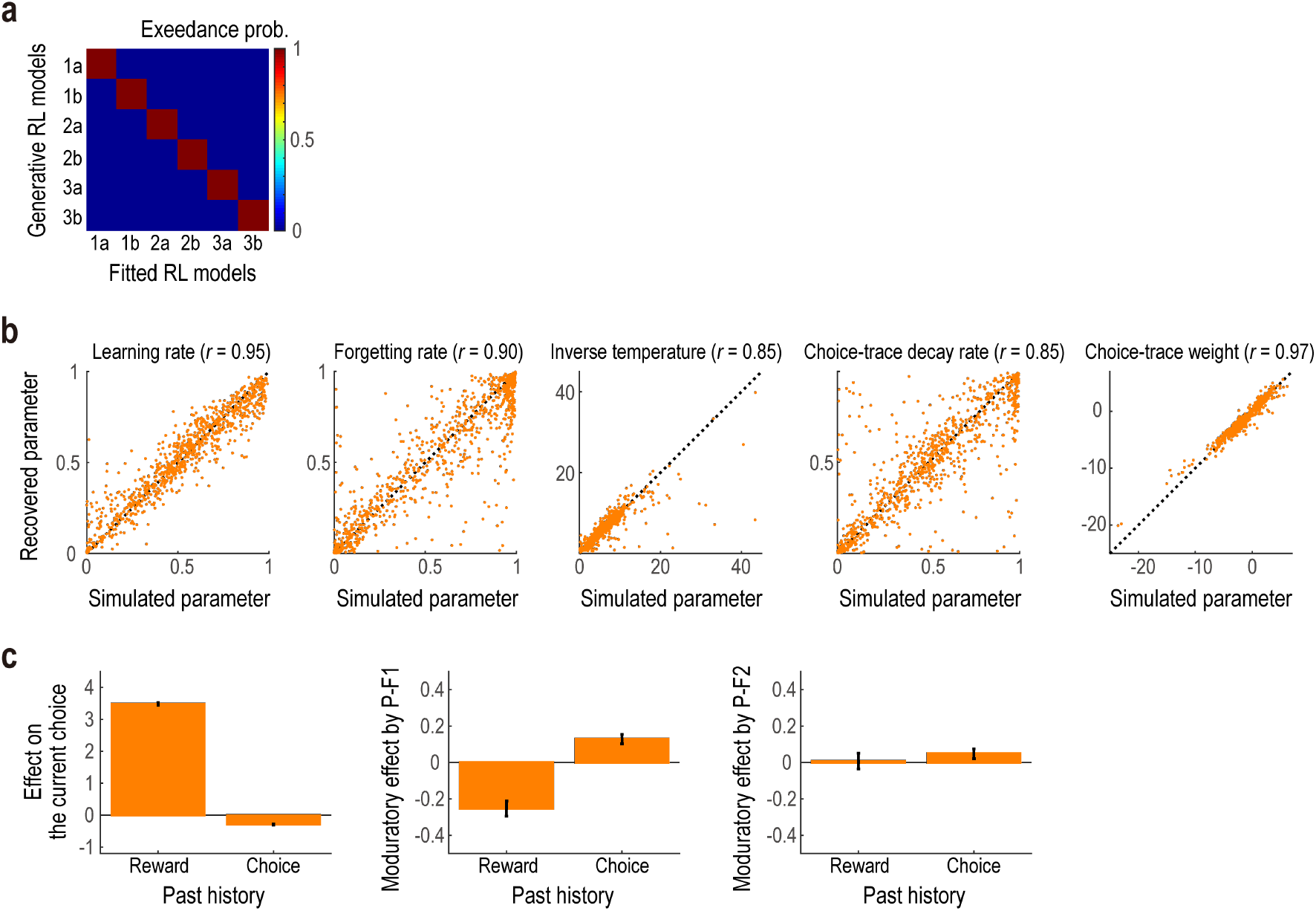
Validation of the computational models and the model-fitting procedure based on simulation data. (a) Confusion matrix to evaluate the performance of our Bayesian model selection. Each row denotes exceedance probabilities of the six competing models (higher values close to *one* indicate a better fit) on the data generated by the corresponding model. For example, the first row depicts exceedance probabilities of the six models on the data generated by the first model (i.e., RL1a). The diagonal elements are found to be close to *one*, meaning that each of these models is identifiable by the model selection procedure. (b) Parameter recovery analysis. We simulated the data based on the best-fitted model (i.e., RL2b) and the parameters, and then try to recover the parameter values by fitting the model to the simulated data. For each of the five decision parameters, the recovered parameter estimates are plotted against the original ones used in the simulation. (c) Replication of the model-neutral regression analysis on the simulated data. Here, we checked whether the results described in Fig. 3 are successfully replicated on the simulated data. The format is the same as in Fig. 3.

**Fig. S3:**
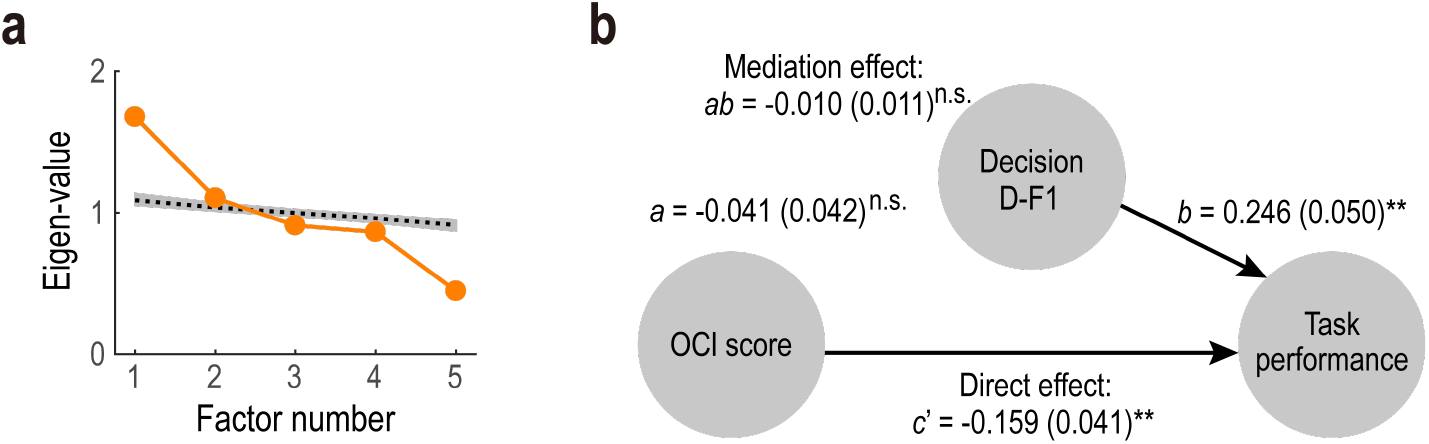
Supplementary information about the computational model-based analyses. (a) Scree-plot of the decision parameters in the best-fitted model (RL2b). We plot eigen values derived from the cross-correlation matrix (Fig. 4b) in *orange*. The black dashed line and the shaded area denote the results of the parallel analysis: that is, eigen values derived from the random data (median, 5% and 95% quantiles). (b) Mediation path diagram for the OCI score, the first decision factor (D-F1) and the task performance (*n* = 939). OCI, Obsessive-Compulsive Inventory. The format is the same as in Fig. 4e. \*\**P* < 0.01; and n.s., non-significant. Path *a*: *P* = 0.284, *Bootstrap* test; path *b*: *P* = 0.003, *Bootstrap* test; path *ab*: *P* = 0.305, Bootstrap test; and path *c*: *P* < 0.001, *Bootstrap* test.

## References

1. Rangel, A., Camerer, C. & Montague, R. P. A framework for studying the neurobiology of value-based decision making. Nat Rev Neurosci 9, 545–556 (2008).

2. Endrass, T., Koehne, S., Riesel, A. & Kathmann, N. Neural correlates of feedback processing in obsessive–compulsive disorder. J Abnorm Psychol 122, 387 (2013).

3. Nielen, M., den Boer, J. & Smid, H. Patients with obsessive–compulsive disorder are impaired in associative learning based on external feedback. Psychol Med 39, 1519–1526 (2009).

4. Chen, C., Takahashi, T., Nakagawa, S., Inoue, T. & Kusumi, I. Reinforcement learning in depression: A review of computational research. Neurosci Biobehav Rev 55, 247–267 (2015).

5. Mkrtchian, A., Aylward, J., Dayan, P., Roiser, J. P. & Robinson, O. J. Modeling Avoidance in Mood and Anxiety Disorders Using Reinforcement Learning. Biol Psychiat 82, 532–539 (2017).

6. Strauss, G. P., Waltz, J. A. & Gold, J. M. A Review of Reward Processing and Motivational Impairment in Schizophrenia. Schizophrenia Bull 40, S107–S116 (2014).

7. Sutton, R. S. & Barto, A. G. Reinforcement Learning: An Introduction. (1998).

8. Wilson, R. C., Geana, A., White, J. M., Ludvig, E. A. & Cohen, J. D. Humans use directed and random exploration to solve the explore–exploit dilemma. J Exp Psychology Gen 143, 2074 (2014).

9. Schulz, E. & Gershman, S. J. The algorithmic architecture of exploration in the human brain. Curr Opin Neurobiol 55, 7–14 (2019).

10. Song, M., Bnaya, Z. & Ma, W. Sources of suboptimality in a minimalistic explore–exploit task. Nat Hum Behav 3, 361–368 (2019).

11. Daw, N. D., O’Doherty, J. P., Dayan, P., Seymour, B. & Dolan, R. J. Cortical substrates for exploratory decisions in humans. Nature 441, 876 (2006).

12. Frank, M. J., Doll, B. B., Oas-Terpstra, J. & Moreno, F. Prefrontal and striatal dopaminergic genes predict individual differences in exploration and exploitation. Nat Neurosci 12, nn.2342 (2009).

13. Schultz, W., Dayan, P. & Montague, R. P. A Neural Substrate of Prediction and Reward. Science 275, 1593–1599 (1997).

14. O’Doherty, J. et al. Dissociable Roles of Ventral and Dorsal Striatum in Instrumental Conditioning. Science 304, 452–454 (2004).

15. Camerer, C. & Ho, T. Experience-weighted Attraction Learning in Normal Form Games. Econometrica 67, 827–874 (1999).

16. Erev, I. & Roth, A. E. Predicting How People Play Games: Reinforcement Learning in Experimental Games with Unique, Mixed Strategy Equilibria. The American Economic Review 88, 848–881

17. Rescorla, R. A. & Wagner, A. R. A theory of Pavlovian conditioning: Variations in the effectiveness of reinforcement and nonreinforcement. 64–99

18. Rutledge, R. B., Dean, M., Caplin, A. & Glimcher, P. W. Testing the Reward Prediction Error Hypothesis with an Axiomatic Model. J Neurosci 30, 13525–13536 (2010).

19. Suzuki, S. et al. Learning to Simulate Others’ Decisions. Neuron 74, 1125–1137 (2012).

20. Lee, D. Decision Making: From Neuroscience to Psychiatry. Neuron 78, 233–248 (2013).

21. Montague, R. P., Dolan, R. J., Friston, K. J. & Dayan, P. Computational psychiatry. Trends Cogn Sci 16, 72–80 (2012).

22. Maia, T. V. & Frank, M. J. From reinforcement learning models to psychiatric and neurological disorders. Nat Neurosci 14, 154 (2011).

23. Huys, Q. J., Maia, T. V. & Frank, M. J. Computational psychiatry as a bridge from neuroscience to clinical applications. Nat Neurosci 19, 404–413 (2016).

24. Gillan, C. & Robbins, T. Goal-directed learning and obsessive-compulsive disorder. Philosophical Transactions Royal Soc B Biological Sci 369, 20130475–20130475 (2014).

25. Gillan, C. M. et al. Disruption in the Balance Between Goal-Directed Behavior and Habit Learning in Obsessive-Compulsive Disorder. Am J Psychiatry 168, 718–726 (2011).

26. Gillan, C. M., Kosinski, M., Whelan, R., Phelps, E. A. & Daw, N. D. Characterizing a psychiatric symptom dimension related to deficits in goal-directed control. Elife 5, e11305 (2016).

27. Huys, Q. J., Pizzagalli, D. A., Bogdan, R. & Dayan, P. Mapping anhedonia onto reinforcement learning: a behavioural meta-analysis. Biology Mood Anxiety Disord 3, 12 (2013).

28. Kunisato, Y. et al. Effects of depression on reward-based decision making and variability of action in probabilistic learning. J Behav Ther Exp Psy 43, 1088–1094 (2012).

29. Strauss, G. P. et al. Deficits in Positive Reinforcement Learning and Uncertainty-Driven Exploration Are Associated with Distinct Aspects of Negative Symptoms in Schizophrenia. Biol Psychiat 69, 424–431 (2011).

30. Harlé, K. M., Guo, D., Zhang, S., Paulus, M. P. & Yu, A. J. Anhedonia and anxiety underlying depressive symptomatology have distinct effects on reward-based decision-making. Plos One 12, e0186473 (2017).

31. Caspi, A. et al. The p Factor. Clin Psychological Sci 2, 119–137 (2013).

32. Newman, D. L., Moffitt, T. E., Caspi, A. & Silva, P. A. Comorbid mental disorders: Implications for treatment and sample selection. J Abnorm Psychol 107, 305 (1998).

33. Redish, D. A. & Gordon, J. A. Computational Psychiatry. (2016).

34. Feczko, E. et al. The Heterogeneity Problem: Approaches to Identify Psychiatric Subtypes. Trends Cogn Sci (2019). doi:10.1016/j.tics.2019.03.009

35. Gillan, C. M. & Daw, N. D. Taking Psychiatry Research Online. Neuron 91, 19–23 (2016).

36. Akaishi, R., Umeda, K., Nagase, A. & Sakai, K. Autonomous Mechanism of Internal Choice Estimate Underlies Decision Inertia. Neuron 81, 195–206 (2014).

37. Gershman, S. J., Pesaran, B. & Daw, N. D. Human Reinforcement Learning Subdivides Structured Action Spaces by Learning Effector-Specific Values. J Neurosci 29, 13524–13531 (2009).

38. Lau, B. & Glimcher, P. W. DYNAMIC RESPONSE-BY-RESPONSE MODELS OF MATCHING BEHAVIOR IN RHESUS MONKEYS. J Exp Anal Behav 84, 555–579 (2005).

39. Walton, M. E., Behrens, T., Buckley, M. J., Rudebeck, P. H. & Rushworth, M. Separable Learning Systems in the Macaque Brain and the Role of Orbitofrontal Cortex in Contingent Learning. Neuron 65, 927–939 (2010).

40. Katahira, K. The relation between reinforcement learning parameters and the influence of reinforcement history on choice behavior. J Math Psychol 66, 59–69 (2015).

41. Ito, M. & Doya, K. Validation of Decision-Making Models and Analysis of Decision Variables in the Rat Basal Ganglia. J Neurosci 29, 9861–9874 (2009).

42. Katahira, K., Yuki, S. & Okanoya, K. Model-based estimation of subjective values using choice tasks with probabilistic feedback. J Math Psychol 79, 29–43 (2017).

43. Toyama, A., Katahira, K. & Ohira, H. A simple computational algorithm of model-based choice preference. Cognitive Affect Behav Neurosci 17, 764–783 (2017).

44. Kato, A. & Morita, K. Forgetting in Reinforcement Learning Links Sustained Dopamine Signals to Motivation. Plos Comput Biol 12, e1005145 (2016).

45. Katahira, K. The statistical structures of reinforcement learning with asymmetric value updates. J Math Psychol 87, 31–45 (2018).

46. Palminteri, S., Lefebvre, G., Kilford, E. J. & Blakemore, S.-J. Confirmation bias in human reinforcement learning: Evidence from counterfactual feedback processing. Plos Comput Biol 13, e1005684 (2017).

47. Stephan, K., Penny, W. D., Daunizeau, J., Moran, R. J. & Friston, K. J. Bayesian model selection for group studies. Neuroimage 46, 1004–1017 (2009).

48. Wilson, R. C. & Collins, A. Ten simple rules for the computational modeling of behavioral data. doi:10.31234/osf.io/46mbn

49. Wager, T. D., Davidson, M. L., Hughes, B. L., Lindquist, M. A. & Ochsner, K. N. Prefrontal-Subcortical Pathways Mediating Successful Emotion Regulation. Neuron 59, 1037–1050 (2008).

50. Atlas, L. Y., Bolger, N., Lindquist, M. A. & Wager, T. D. Brain Mediators of Predictive Cue Effects on Perceived Pain. J Neurosci 30, 12964–12977 (2010).

51. Ishikawa, R., Kobori, O. & Shimizu, E. Development and validation of the Japanese version of the obsessive-compulsive inventory. Bmc Res Notes 7, 306 (2014).

52. Robbins, T. W., Vaghi, M. M. & Banca, P. Obsessive-Compulsive Disorder: Puzzles and Prospects. Neuron 102, 27–47 (2019).

53. Association, A. Diagnostic and Statistical Manual of Mental Disorders (DSM5). (2013).

54. Stewart, N., Chandler, J. & Paolacci, G. Crowdsourcing Samples in Cognitive Science. Trends Cogn Sci 21, 736–748 (2017).

55. Vaghi, M. M. et al. Compulsivity Reveals a Novel Dissociation between Action and Confidence. Neuron 96, 348–354.e4 (2017).

56. Menzies, L. et al. Neurocognitive endophenotypes of obsessive-compulsive disorder. Brain 130, 3223–3236 (2007).

57. Remijnse, P. L. et al. Reduced Orbitofrontal-Striatal Activity on a Reversal Learning Task in Obsessive-Compulsive Disorder. Arch Gen Psychiat 63, 1225–1236 (2006).

58. Lawson, R. P., Mathys, C. & Rees, G. Adults with autism overestimate the volatility of the sensory environment. Nat Neurosci 20, 1293–1299 (2017).

59. Browning, M., Behrens, T. E., Jocham, G., O’Reilly, J. X. & Bishop, S. J. Anxious individuals have difficulty learning the causal statistics of aversive environments. Nat Neurosci 18, 590–596 (2015).

60. Tomov, M., Truong, V., Hundia, R. & Gershman, S. Dissociable Neural Correlates of Uncertainty Underlie Different Exploration Strategies. Biorxiv 478131 (2019). doi:10.1101/478131

61. Maniaci, M. R. & Rogge, R. D. Caring about carelessness: Participant inattention and its effects on research. J Res Pers 48, 61–83 (2014).

62. Raine, A. & Benishay, D. The SPQ-B: A Brief Screening Instrument for Schizotypal Personality Disorder. J Pers Disord 9, 346–355 (1995).

63. Ito, S., Obu, S., Ota, M., Takao, T. & Sakamoto, S. Reliability and validity of the Japanese version of SPQ-B (Schizotypal Personality Questionnaire Brief). Jpn. Bull. Soc. Psychiatry 168–176 (2008).

64. Zung, W. W. A Self-Rating Depression Scale. Arch Gen Psychiat 12, 63–70 (1965).

65. Spielberger, C., Gorsuch, R. & Lushene, R. Manual for the State-Trait Anxiety Inventory (Self-Evaluation Questionnaire). (1970).

66. Patton, J. H., Stanford, M. S. & Barratt, E. S. Factor structure of the barratt impulsiveness scale. J Clin Psychol 51, 768–774 (1995).

67. Yanagisawa, K. et al. Family socioeconomic status modulates the coping-related neural response of offspring. Soc Cogn Affect Neur 8, 617–622 (2013).

68. Noonan, M. P., Chau, B., Rushworth, M. & Fellows, L. K. Contrasting Effects of Medial and Lateral Orbitofrontal Cortex Lesions on Credit Assignment and Decision-Making in Humans. J Neurosci 37, 7023–7035 (2017).

69. Leo, W. R. Techniques for Nuclear and Particle Physics Experiments. (1994).

70. Daw, N. D. Trial-by-trial data analysis using computational models. (2011).

71. Niv, Y., Edlund, J. A., Dayan, P. & O’Doherty, J. P. Neural Prediction Errors Reveal a Risk-Sensitive Reinforcement-Learning Process in the Human Brain. J Neurosci 32, 551–562 (2012).

72. West, S. G., Finch, J. F. & Curran, P. J. Structural equation models with nonnormal variables: Problems and remedies. (1995).

73. Fabrigar, L. R., Wegener, D. T., MacCallum, R. C. & Strahan, E. J. Evaluating the use of exploratory factor analysis in psychological research. Psychol Methods 4, 272 (1999).

